# Design of extended metal-binding β-sandwiches from *de novo* immunoglobulin domains

**DOI:** 10.1101/2022.08.03.502643

**Authors:** Jorge Roel-Touris, Marta Nadal, Enrique Marcos

**Affiliations:** Protein Design and Modeling Lab, Department of Structural Biology, Molecular Biology Institute of Barcelona (IBMB-CSIC), Baldiri Reixac 15, 08028 Barcelona, Spain

## Abstract

The antigen binding sites of antibodies, and derivatives such as scFvs, consist of an immunoglobulin structural framework that anchors hypervariable loops, and that is arranged as a dimer of β-sandwiches (from the heavy and light chains) packing face-to-face. Yet, the naturally occurring dimer orientation of antibodies is not well suited for engineering rigid single-chain formats of increased stability and solubility, which are two key limitations of engineered antibodies. In this work, we computationally designed an extended 12-stranded β-sandwich as a rigid single-chain dimer of *de novo* immunoglobulin domains interacting through an alternative edge-to-edge arrangement. The extended β-sandwich was found to be hyperstable and structurally accurate as confirmed through X-ray crystallography. We functionalized this design by inserting a long EF-hand calcium-binding motif into the β-hairpin bridge between both domains, showing high metal-binding affinity while remaining soluble and stable in solution. Finally, we propose a few designs with two EF-hand motifs in either or the same side of the scaffold with high-confidence structure predictions; altogether suggesting the robustness and versatility of our scaffold to harbor long functional loops. Our extended β-sandwiches are structurally divergent to natural antibodies and open new avenues for incorporating multiple loop binding sites either for increased or bispecific activities.

## INTRODUCTION

Engineered antibodies have revolutionized the treatment of cancer and auto-immune diseases, becoming nowadays the largest growing class of pharmaceutical drugs (*1, 2*). Several alternative formats have been engineered to address some recurrent limitations of natural antibodies (*3–5*). One of these formats are single-chain variable fragments (scFvs), which are antibody-based constructs connecting the variable regions of the heavy (VH) and light (VL) chains of natural antibodies with flexible linkers (*6*). These simplified structures have several advantages over full-length antibodies since they are smaller in size, suitable for recombinant expression in bacteria and, as a single polypeptide, amenable to display optimization technologies. However, the dependence of engineered antibodies on the naturally occurring structure of the immunoglobulin-like (Ig) domain components and their interfaces, poses several limitations. The scFv linker needs to be quite long (10-25 amino acids) due to the large distance separation between the N and C termini of the two subunits. This condition can compromise stability and structural control, as seen in many other fusion constructs combining different protein domains (*7*). Additionally, the dimer interface between the VH and VL Ig domains is formed by two β-sheets packing face-to-face, and in such arrangement edge β-strands are solvent-exposed and can form edge-to-edge intermolecular interactions – i.e. through backbone hydrogen bonding involving the exposed amine and carbonyl groups – and therefore increase aggregation propensity (*8*). The ability of custom-designing Ig structures and interfaces not tied to natural ones should allow, for example, to generate more rigid and stable scFv-like formats.

*De novo* designing Ig domains with new structures and dimer interfaces could give rise to alternative antibody-like formats with superior properties. We have recently described principles for designing single-domain Ig proteins *de novo* (*9*). Among these *de novo* Ig proteins (dIGs), one design was found to be dimeric and thermostable in solution (dIG14). It was originally designed as a monomeric 7-stranded immunoglobulin, but the crystal structure revealed a homodimer with an edge-to-edge interface forming two extended β-sheets, overall building a 12-stranded (6×6) β-sandwich (Fig. 1). Such serendipitous structure is constrained by the requirement of backbone hydrogen bond pairing between two pairs of edge β-strands from four β-sheets simultaneously, and not all Ig domain structures are compatible with such arrangement. The high number of backbone hydrogen bonds at this interface contributes to dimer stability, and their distribution in two layers increases structural rigidity. From the design perspective, harnessing the strength of edge-to-edge β-strand interactions in a two-layer dimer interface can be a powerful strategy for stabilizing and rigidifying Ig dimers, while favoring solubility (as this reduces the number of exposed β-strand edges). Here, we set out to computationally redesign the dIG14 structure as a rigid single-chain Ig dimer (dIG-scdim) bridging the two domains with structured loops. We found that the dIG14 single-chain dimer can be designed with very short loop connections, exceptional stability, and structural accuracy as confirmed by X-ray crystallography. We also found that the bridge between monomers is a well-suited spot for scaffolding EF-hand calcium-binding loops; overall suggesting the robustness of the structure and potential for harboring multiple functional sites against a broader range of targets.

**Fig. 1.**
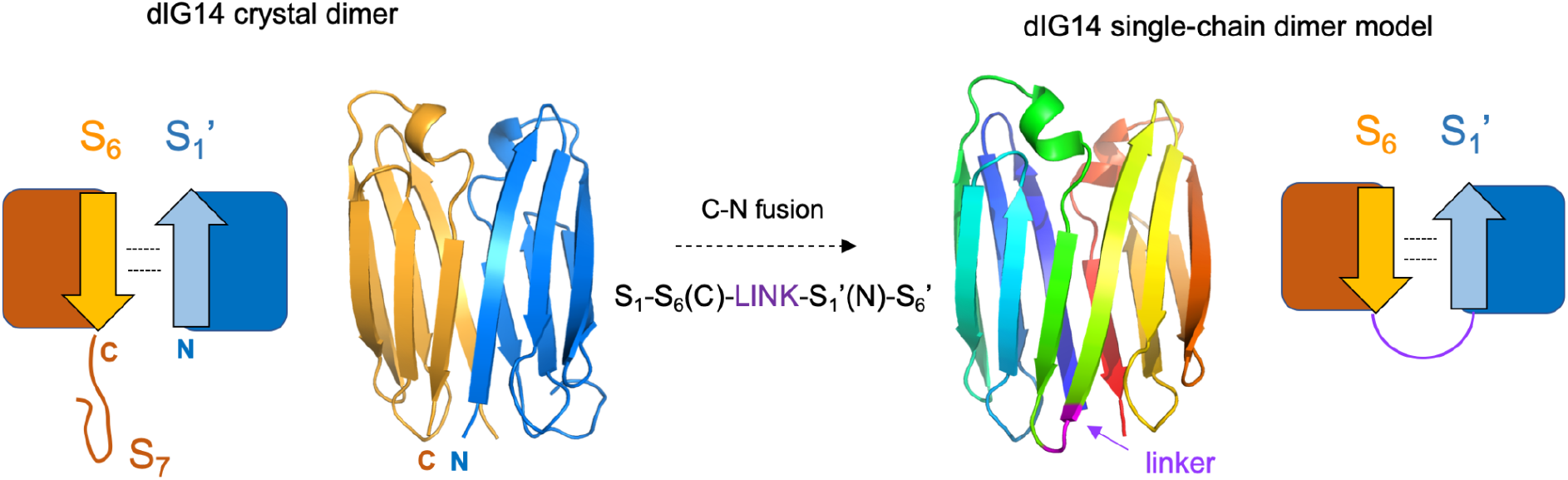
The *de novo* designed immunoglobulin dIG14 as a single-chain dimer. Fusing the two 6-stranded Ig domains involved in the dIG14 dimer interface (*left*) with a short linker enables formation of a 12-stranded single-chain Ig dimer. The dIG14-scdim design model (*right*) was built by loop insertion between the two chains of the crystal structure after removing the C-terminal β-strand.

## RESULTS

### Extended β-sandwiches based on dIG14

The dIG14 structure was effectively formed by two 6-stranded Ig monomers, suggesting that the C-terminal β-strand was dispensable for proper folding. As the sixth β-strand of one monomer and the first β-strand of the second build an antiparallel interface (and therefore orient their C- and N-termini in close proximity), we reasoned that the two Ig interacting monomers could be fused with a short linker forming a β-hairpin at the interface. (Fig. 1). Based on the dIG14 crystal structure, we removed the last 9 residues of the sequence, and to find the shortest connection between the two chains we performed Rosetta (*10*) fragment-based insertion of poly-glycine loops (ranging between 2 and 5 amino acids) connecting W68 or G69 of one monomer with G1 or R2 of the other monomer. We found that the gap could be easily bridged with minimal backbone strain with loops equal or larger than 2 connecting G69 with G1. For these single-chain dimers, AlphaFold2 (*11*) (AF2) generated highly confident predictions (pLDDT > 90) across all residue positions and that matched very closely the design model (Cα-RMSD = 0.6 Å) (Fig. 2a).

**Fig. 2.**
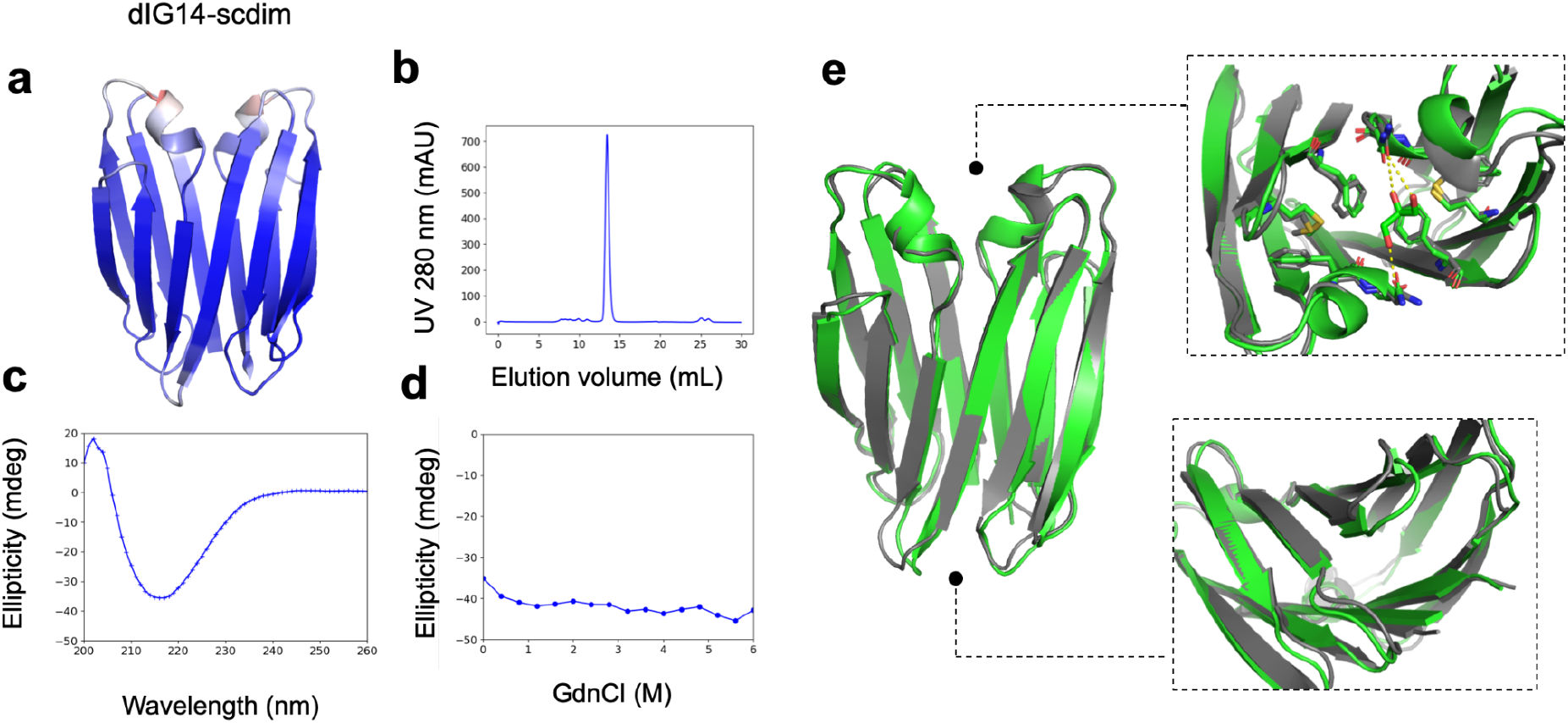
Design and experimental characterization of the dIG14 single-chain dimer. **a**, First AlphaFold2 model of the single-chain dimer formed by a short GG linker, and colored by pLDDT (from red to blue increasing in pLDDT; scale 70 to 100). **b**, Size-exclusion chromatogram of the expressed and purified single-chain dimer, which is monodisperse and elutes at 13.4 mL as expected for a protein in this size range. **c**, Far-ultraviolet circular dichroism spectra at 25 °C. **d**, Chemical denaturation with GdnCl monitored with circular dichroism at 215 nm and 25 °C. **e**, dIG14-scdim design model (*green*) in comparison with the crystal structure (*gray*). Right insets show *top* and *bottom* views of the structural agreement across the two β-sheets. A small ligand-binding cavity is formed at the top *via* hydrogen bond and aromatic interactions.

Encouraged by the confident predictions, we selected the dIG14-scdim designed with a GG linker for experimental characterization, and ordered a synthetic gene encoding for the designed sequence. We express it in *Escherichia coli*, purified it by affinity and size-exclusion chromatography, and it was found to be well-expressed, soluble and monomeric by size-exclusion chromatography combined with multi-angle light-scattering (SEC-MALS) (Fig. 2b). Moreover, it was found to have far-UV circular dichroism spectra characteristic of all-β proteins and turned out to be hyperstable by circular dichroism (Fig. 2c, d) – the protein remains folded in 6 M GdnCl. We succeeded in solving a crystal structure of dIG14-scdim at 2.8 Å resolution (Table S1), and was found in excellent agreement with the computational model across the 12 β-strands (Cα-RMSD = 0.8 Å; Fig. 2e). This structure can be regarded as a flattened β-barrel. At the bottom, the designed linker is surrounded by a tightly packed area stabilized by aromatic stacking, and at the top it was found a cavity binding a glycerol molecule (as crystallization component) that is surrounded by the two β-arch helices (Fig. 2e) – the structure of this area thus could be also diversified for designing small-molecule binding sites. The complex β-sheet arrangement of the structure constitutes a novel all-β domain topology, given that the closest structural analogues found in the PDB or the AlphaFold Structure Database (*12*) had low TM-scores (*13*) (<=0.65), and those were β-sandwiches formed by a different number of β-strands and strand pairing organization.

### Computational design and characterization of functional loops

We reasoned that the β-hairpin bridge between monomer subunits, besides playing a structural role, could also be well-suited for grafting functional loops. More generally, β-hairpin loop regions are in principle better suited for loop grafting as compared to β-arch loop regions, which tend to fold more slowly and hence need to be more structured (*14*). We computationally grafted an EF-hand calcium-binding motif (PDB accession code 1NKF(*15*)) in the loop connecting both subunits (G70-G71), and designed N- and C-terminal linkers as short and structured as possible to stabilize the motif into our structure using RosettaRemodel (*16*). We generated 772 designs with grafted EF-hand motifs containing between 15 and 21 amino acids, and subsequently analyzed them with AF2. We first probed protein folding by checking whether AF2 predictions closely matched the structure of the β-sandwich scaffold (i.e. not considering the binding motif in the analysis) to the parent dIG14-scdim design. More than one third of the grafted designs had at least one (out of five) high-confidence AF2 prediction (pLDDT > 90) with a Cα-RMSD < 1 Å to the scaffold. Furthermore, 80 designs had at least 3 high-confidence AF2 predictions with scaffold Cα-RMSDs < 1 Å, and for 22 of them, all 5 predictions were within these ranges; suggesting the robustness and potential of the dIG14-scdim structure to scaffolding functional loops.

We identified four designs with high-confidence AF2 predictions closely recapitulating the design structure, both at the scaffold and binding motif levels (0.66 Å < total RMSDs < 2.68 Å and 77.96 < pLDDT < 96.3). For the best performing model, *model062_2_EF_1*, we found RMSDs < 1 Å and pLDDTs > 95 (Table S2). Note here the size and folding complexity of the protein scaffold in combination with inserted functional motifs containing between 15 and 21 amino acids. This is a very stringent design validation test considering that the predictions are performed in the absence of the ligand.

The structural deviations shown by the AlphaFold2 predictions are higher on the EF-hand motifs; possibly indicating certain flexibility on the inserted constructs. To investigate on the structural rigidity of the binding motifs, we ran molecular dynamics simulations for the 4 selected designs in the absence of the ligand (Fig. 3b). We computed Root Mean Square fluctuations (RMSFs) for each residue positions and found that the scaffold (except for the β-arch loop regions) remains very rigid (with averaged fluctuations no greater than 1.5 Å), but the motif region seems much more flexible, having RMSF values up to 4 Å. Among the 4 simulated designs, *model018_2_EF_7* shows the highest flexibility on the motif and samples the most divergent loop conformations as compared to the EF-hand native loop (Cα-RMSDs = 3.27 ± 0.98 Å). A previous study, scaffolding similar EF-hand motifs, stressed the importance of the stability of such loops for ligand binding purposes (*17*). Indeed, analyses of the mode of action revealed that the first lysine from the motif acts as an anchor by making hydrogen bonds. This specific contact stabilizes the construct and facilitates the binding of the ion; a fact confirmed by the crystal structure (PDB accession code 6OHH). Similarly, in all of our selected designs, the first lysine of the motif makes hydrogen bonds with either an aspartic or glutamic acid present on the N-terminal linkers. Compelled by these findings, we sought to investigate whether this salt bridge has any effect on the flexibility of the motif, and monitored the pairwise side chain center-of-mass distances between the lysine and either acidic residue throughout the simulation. For the first 3 designs, the pairwise distances between the salt-bridging residues remain within a close interaction distance throughout the simulation, but for *model018_2_EF_7* the contact is lost. Interestingly, these results seem to correlate with the overall flexibility of the motifs and suggest that this specific contact is key in the stability of the grafted EF-hand metal-binding loop into our scaffold.

**Fig. 3.**
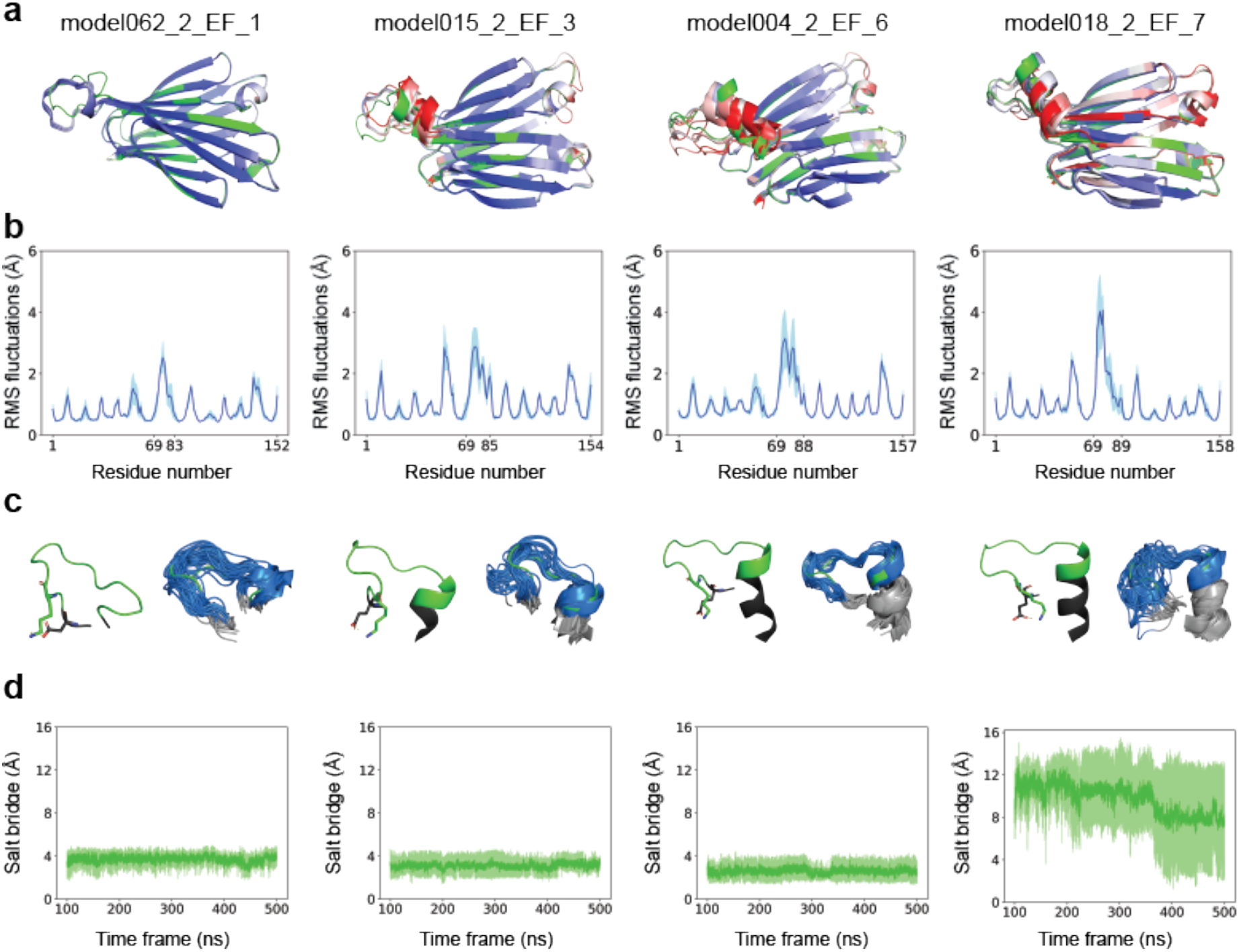
Computational characterization of functional loop scaffolding. **a**, Best performing functionalized designs (green) and their AlphaFold2 predictions colored by pLDDT (from red to blue increasing in pLDDT; scale 70 to 100). **b**, Averaged root mean square fluctuations (RMSFs). Y-axis indicates fluctuations in Angstroms for each residue. X-axis shows the first and last residues of the design and the grafted motif (central region). **c**, Representation of the grafted motifs (left). Ensemble of 30 conformations (10 per replicate) sampled during the molecular dynamics simulations every 50 ns (right). Black/Gray: Designed linkers. Green/Blue: EF-hand metal binding loop. Sticks are shown for the salt-bridging residues. **d**,Pairwise distances between the salt bridge forming residues in Angstroms (y-axis) as a matter of simulation time (x-axis).

### Experimental characterization of a metal-binding design

We selected *model062_2_EF_1* (Fig. 4a) for experimental characterization as it was the design with AlphaFold predictions of highest confidence (pLDDT > 90 for the five models and across all residue positions, including the EF-hand motif) and more similar to the design (Ca-RMSD < 1 Å) (Table S2). For *model062_2_EF_1*, we also noted that the structural rigidity of the binding motif increased in the presence of calcium by molecular dynamics simulations (Fig. 4b), and that the interactions provided by the four EF-hand binding residues (D71, D75 and E82 through their acidic sidechain, and Y77 through the backbone oxygen atom) remained nearly constant throughout the simulation (Fig. 4c). We ordered a synthetic gene encoding for *model062_2_EF_1*, and the designed protein was found to be well-expressed in *E. Coli* and monodisperse by size-exclusion chromatography (Fig. 4d) – eluting slightly before the parent dIG14-scdim, as expected given that it incorporates a 15-residue loop insertion. We next carried out time-resolved terbium luminescence experiments to determine the metal binding ability of the design. Besides calcium, EF-hand motifs can also bind terbium, which has a similar ionic radius and coordination sphere. Terbium provides a convenient binding readout, as its fluorescence can be sensitized by energy transfer from proximal aromatic residues (e.g. tyrosine or tryptophan) when excited at 280 nm. The design was found to bind terbium in a concentration-dependent manner and with a characteristic terbium fluorescence intensity peak at 544 nm (Fig. 4e). Terbium-binding titrations were monitored at 544 nm with *model062_2_EF_1* at 10 μM final concentration, and showed a hyperbolic increase in luminescence that was fitted to a 1:1 binding model, which gave an estimated Kd of 2.8 μM (Fig. 4f). To probe calcium binding, we performed a competitive binding assay by titrating calcium and monitoring the decrease in the terbium fluorescence (of *model062_2_EF_1* mixed with terbium at 10 μM). Calcium was found to compete with terbium for the same binding site (Fig. 4g), but with much lower affinity (Kd = 1.3 mM) - this is expected given the lower charge density of calcium compared to terbium and hence weaker electrostatic interactions with the EF-hand motif.

**Fig 4.**
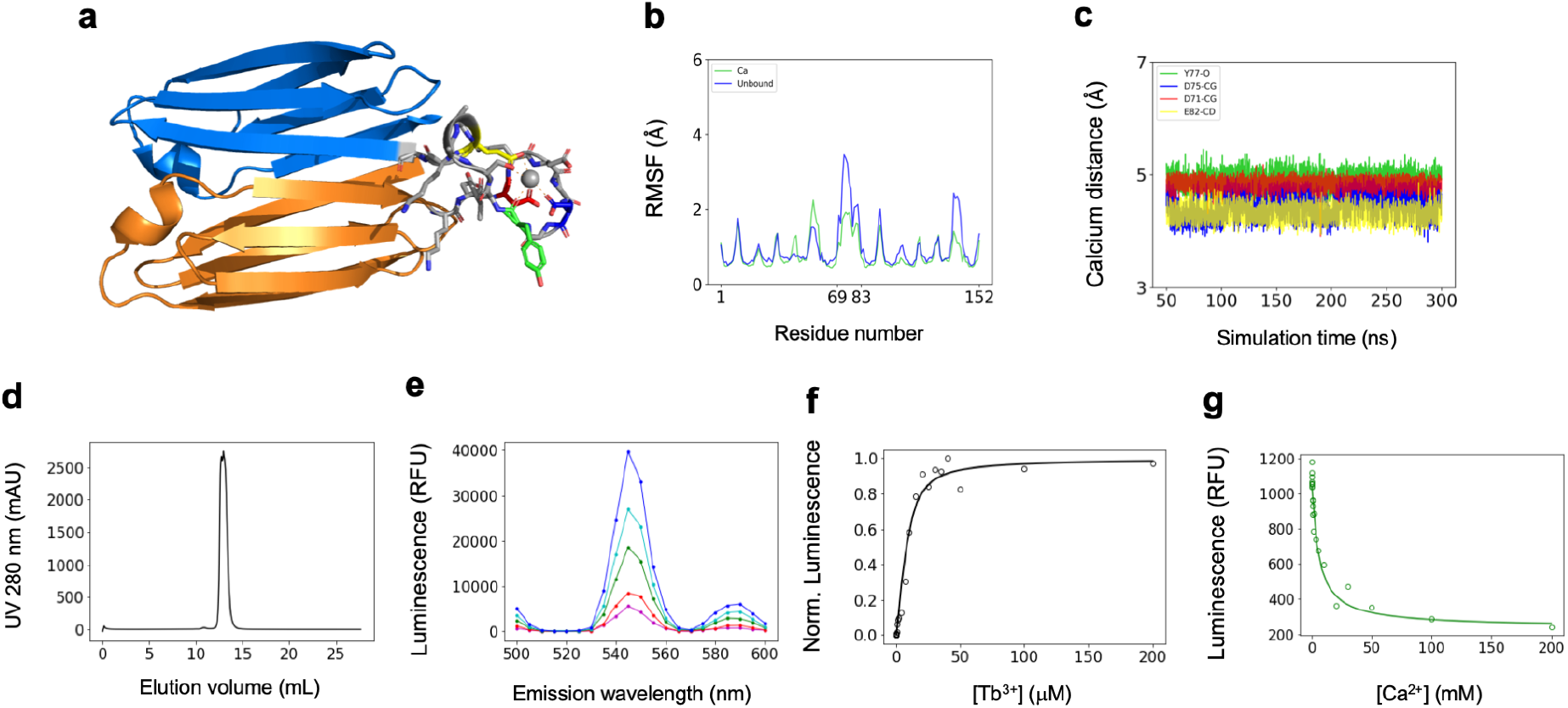
Metal-binding design and experimental characterization. **a**, Computational model of *model062_2_EF_1* with a grafted EF-hand motif (gray) showing Tb^3+^ (gray sphere) bound to EF-hand motif chelating residues (sticks). The two 6-stranded immunoglobulin domains are colored in orange and blue, respectively. EF-hand calcium binding residues are D71 (red), D75 (blue), Y77 (green), E82 (yellow). **b**, Root-mean-square fluctuations (RMSF) obtained from molecular dynamics simulations in the presence and absence of calcium. **c**, Distances between EF-hand motif chelating residues and calcium monitored by molecular dynamics simulations. The four series match the colors of the four binding residues as in (a). **d**, Size-exclusion chromatogram of *model062_2_EF_1*, which is monodisperse and elutes at 12.9 mL. **e**, Time-resolved luminescence emission spectra in different Tb^3+^ final concentrations (magenta: 1.7 μM; red: 3.5 μM; green: 7 μM; cyan: 10 μM; blue: 20 μM) and *model062_2_EF_1* at 10 μM. Time-resolved luminescence intensity is given in relative fluorescence units (RFU). **f**, Tb^3+^ concentration-dependent time-resolved luminescence intensity of 10 μM *model062_2_EF_1* using excitation wavelength λex = 280 nm and emission wavelength λem = 544 nm. Normalized intensities are fit to a one-site binding model by non-linear least squares regression (Kd = 2.8 μM). **g**, Time-resolved luminescence intensity in relative fluorescence units (RFU) for Ca^2+^ titrations with 10 μM *model062_2_EF_1* and 10 μM Tb^3+^, showing Ca^2+^ competition with Tb^3+^ for the *model062_2_EF_1* binding site.

## DISCUSSION

We have designed, characterized and functionalized a 12-stranded β-sandwich from *de novo* immunoglobulin domains. This work builds upon our previous study, where we developed and applied design rules to *de novo* design stable antibody-like scaffolds, subsequently confirmed by X-ray crystallography (*9*). For one of such immunoglobulins (dIG14), originally designed as a monomeric 7-stranded scaffold, X-ray studies revealed a dimer formed by two antiparallel 6-stranded Ig monomers instead; suggesting a route to the design of edge-to-edge single-chain immunoglobulin dimers. Inspired by this discovery, we redesigned our original 7-stranded scaffold into a 6-stranded domain and connected the two domains with the shortest linker possible to effectively create a rigid single-chain immunoglobulin-like dimer (dIG14-scdim). Despite the high folding complexity of the resulting protein topology, which is not observed in nature, this reimagination showed accurate and confident structure predictions, and experimentally was found to be monomeric, extremely stable, and its structure was confirmed by X-ray crystallography. We show that the bridge between the two immunoglobulin-like domains can be readily functionalized by designing a high-affinity metal-binding loop, which suggests that the dIG14-scdim scaffold is robust to long loop insertions.

Loops connecting adjacent β-strands are well-suited spots for scaffolding functional motifs. Functional loops are typically longer and less structured than canonical β-hairpin motifs frequently used in *de novo* proteins, and must therefore be grafted in such a way that the overall stability of the scaffold is not compromised. To this end, we computationally designed N- and C-terminal linkers to accommodate the EF-hand to the scaffold. Among all of our designs, the most promising ones had N-terminal linkers composed of two residues and C-terminal linkers of varying length (from one to seven residues). Interestingly, the N-terminal linkers always had a negatively charged residue (either aspartic or glutamic acid) immediately preceding the EF-hand motif and one residue away from a lysine (second residue of the motif); forming a salt bridge. By molecular dynamics simulations, we noted that this contact confers a more rigid nature to the motif and stabilizes the metal-binding loop; overall resulting in better binding capacities as confirmed by the experimental assays.

A monomeric 6-stranded immunoglobulin scaffold has two β-hairpins connecting strands 1-2 and 3-4 (connections 2-3, 4-5 and 5-6 are β-arches); pointing towards the same side of the scaffold. However, our dIG14-scdim has five β-hairpins connecting strands 1-2 and 3-4 (on one monomer), 7-8 and 9-10 (on the other monomer) and 6-7 as the bridge between both monomers; spanning both sides of the scaffold. These extra β-hairpin motifs open the possibility of scaffolding multiple loops at either or same side; especially interesting for increased or bispecific activity. As an example, we computationally designed and characterized a few scaffolds with two EF-hand motifs (Fig. S1). The metal-binding motifs were grafted at the same or opposite sites, mimicking somehow scaffolds with increased or, if functionally diverse, bispecific activity. The scaffold was found to be robust to multiple loop insertions based on AlphaFold2 structure prediction (Fig. S1 and Table S3).

The single-chain dimer dIG14-scdim falls in a size range (140 amino acids) that is too large for *ab initio* structure prediction (*18*), which has been the gold-standard in silico test for *de novo* designed proteins during the last decade (*19*). AlphaFold2 successfully predicted very accurate models with high confidence for dIG14-scdim and a substantial number of functionalized variants harboring long loop insertions, which represent an even more challenging prediction. The advent of deep-learning structure prediction breaks a historical size limit set by *ab initio* structure prediction, and now allows us to make accurate structural validations for larger proteins than before; opening the door to the *de novo* design of increasingly large and complex folds, including multidomain proteins. By combining accurate structure prediction with molecular dynamics simulations assessing the binding motif structural rigidity, we identified an excellent design construct (*model062_2_EF_1*) straight from computation that was found to be stable in solution and showed high-affinity terbium binding.

Overall, extending the two β-sheets of immunoglobulin domains through edge-to-edge interactions and short or structured connections constitutes an effective design strategy for generating hyperstable scaffolds with multiple positions amenable for inserting binding loop motifs, such as complementarity-determining regions of antibodies or nanobodies. Interdomain interfaces formed by two-layer edge-to-edge strand pairing minimizes the number of exposed β-strand edges prone to self-association, and therefore contributes to favor monomeric stability in solution and solubility. The overall structure of such edge-to-edge single-chain immunoglobulin dimers could be readily diversified by tuning the structure, number of β-strands or topology of the Ig monomers and the orientation of the inter-domain interfaces (via either parallel or antiparallel edge β-strand pairings). Extended β-sandwiches from *de novo* immunoglobulin domains can be particularly useful as robust antibody-like scaffolds with superior biophysical properties and versatility for combining binding motifs.

## MATERIALS AND METHODS

### Design of linkers

We bridged the two domains by designing short loop segments using the Blueprint Builder (*20*) mover implemented in RosettaScripts (*21*). Poly-glycine loop fragments ranging between 2 and 5 residues were inserted between W68 or G69 of one domain and G1 or R2 of the other domain. Fragment insertion was performed using the fldsgn_cen centroid scoring function.

### Design of functional loops

The Protein Data Bank (PDB) accession code 1NKF (*15*) was retrieved and only the 12 metal-binding residues (DKDGDGYISAAE) were kept. Blueprint files were generated for both the dIG14 single-chain dimer and the EF-hand motif. Additionally, 18 blueprint files were created for domain insertion by sampling all combinations of N-terminal linkers between 2 and 3 residues and C-terminal linkers spanning 0 to 8 residues. The N-terminal linkers were forced to be in an extended conformation, while the C-terminal linkers in α-helical conformation. RosettaRemodel (*16*) was run 100 times for each of the domain insertion blueprints while designing the sequence of both linkers; restricted to a series of residues specified in their corresponding blueprint files. The metal non-coordinating residues were allowed to repack while the metal-binding residues were kept fixed. 598 out the 1,800 simulations produced at least one closed structure; resulting in a total of 772 unique EF-hand grafted single-chain models.

### Protein structure prediction

The local installation (LocalColabFold: github.com/YoshitakaMo/localcolabfold) of the optimized AlphaFold2 software version (ColabFold) (*22*) was used for protein structure predictions. The mmseqs2 method (*23*)was chosen to compute the Multiple Sequence Alignment (MSA) with the –use_turbo option enabled for optimal speed. The number of recycles was kept at 3 (default) and the predictions were relaxed using the Amber force field. Several structural metrics from the predictions including the total Root Mean Square Deviation (RMSD-total), RMSD of the scaffold (RMSD-scaffold; excluding the inserted domain), RMSD of each of the linkers (RMSD-linker1/2) and the RMSD of EF-hand binding motif (RMSD-motif) against the grafted model were calculated for filtering purposes. First, all predictions with RMSDs >1 Å (excluding RMSD-total) were discarded. This filter returned 351 predictions out of 3860 (772 grafted models * 5 AF2 predictions). Next, in order to assure prediction convergence, only those grafted models with at least 4 AF2 predictions passing these cutoffs were kept; yielding 8 models in total. Finally, the best 4 designs according to their RMSD-total and pLDDT values were selected for further analysis.

### Molecular dynamics simulations

The 4 selected designs were used as starting points for molecular dynamics simulations. Dodecahedron TIP3P water boxes were employed to solvate the proteins with a buffer distance of 11 Å to the box edges. Na^+^ and Cl^-^ ions were added to provide charge neutrality at a total concentration of 150 mM. The Amber14SB force field (*24, 25*)was used for proteins and NaCl, and Ca^2+^ parameters were obtained from Bradbrook et al (*26*). First, the systems were minimized using the steepest descent method. Next, the systems were equilibrated involving an initial heating to 100 K at constant volume for 50 ps followed by heating to 298 K at a constant pressure of 1 bar. Production runs were performed with GROMACS (*27*) version 2018.3 at 1 bar and 298 K with periodic boundary conditions and a 2 fs timestep, with nonbonded short-range interactions calculated within a cutoff of 10 Å. Each of the 4 designs was simulated in three independent 500 ns replicates. For the simulations including Ca^2+^, each of the two system conditions (absence/presence of Ca^2+^) were simulated in one 300 ns run. The first 100 ns and 50 ns (absence/presence of Ca^2+^) of each production run were discarded from analysis to allow for adequate equilibration from the starting conformation.

### Recombinant expression and purification of the designed proteins

Synthetic genes encoding for the designed amino acid sequences were obtained from Genscript and cloned into the pET-28a-TEV expression vector, with genes inserted within NdeI and XhoI restriction sites and the pET-28a-TEV backbone encoding an N-terminal hexa histidine tag followed by a Tobacco-Etch-Virus peptidase (TEV) cleavage site. *Escherichia coli* BL21 (DE3) competent cells (Sigma) were transformed with these plasmids, and starter cultures from single colonies were grown overnight at 37°C in Luria-Bertani (LB) medium supplemented with kanamycin. Overnight cultures were used to inoculate 500 ml of LB medium supplemented with kanamycin and cells were grown at 37 °C under shaking, until an optical density (OD600) of 0.5-0.7 was reached. Protein expression was induced with 1mM of isopropyl β-D-thiogalactopyranoside (IPTG) and cultures were incubated overnight at 18°C. Cells were harvested by centrifugation and resuspended in 50 mM Tris·HCl, 250 mM sodium chloride, pH 7.5 supplemented with 10 mM imidazole, and EDTA-free cOmplete Protease Inhibitor Cocktail (Roche Life Sciences). Cells were lysed using a cell disrupter (Constant Systems), and soluble protein was clarified by centrifugation. For immobilised-metal affinity chromatography (IMAC), proteins were captured on nickel-sepharose HisTrap HP columns (Cytiva), which had previously been washed and pre-equilibrated with the same buffer plus either 500 mM or 20 mM imidazole, respectively. Column-bound protein was washed with a gradient of 20-to-150 mM imidazole in the protein buffer and eluted with a gradient of 200-to-300 mM imidazole in the same buffer. Proteins were further purified by size-exclusion chromatography using a Superdex 75 10/300 GL (GE Healthcare) column. The expression of purified proteins was assessed by SDS-polyacrylamide gel; and protein concentrations were determined from the absorbance at 280 nm measured on a NanoDrop spectrophotometer (ThermoScientific) with extinction coefficients predicted from the amino acid sequences using the ProtParam tool (https://web.expasy.org/protparam/).

### Circular dichroism

Far-ultraviolet circular dichroism measurements were performed to assess secondary structure content and protein stability using a JASCO spectrometer at CCiTUB. Wavelength scans were measured from 260 to 195 nm using a 1 mm path-length cuvette. Protein samples were prepared in PBS buffer (pH 7.4) at a concentration of 0.3-0.4 mg/mL. Guanidine hydrochloride titrations were performed manually using a 6M GdnCl stock solution dissolved into PBS.

### Size-exclusion chromatography coupled to multiple-angle light scattering (SEC-MALS)

SEC-MALS was performed in a Dawn Helios II apparatus (Wyatt Technologies) coupled to a SEC Superdex 200 Increase 10/3000 column. The column was equilibrated with 30 mM Tris·HCl, 250 mM sodium chloride, pH 8 at 25 °C and operated at a flow rate of 0.5 mL/min. Data processing and analysis proceeded with Astra 7 software (Wyatt Technologies).

### Protein crystallization and structure determination

Crystallization screenings were performed at the joint IRB/IBMB Automated Crystallography Platform. Screening solutions were prepared and dispensed into the reservoir wells of 96×2-well MRC crystallization plates (Innovadyne Technologies) by a Freedom EVO robot (Tecan) using the sitting-drop vapor diffusion method. dIG14-scdim crystals appeared at 20 °C in drops consisting of 100 nL protein solution (at 15 mg/ml in 30 mM Tris pH 8, 250 mM NaCl) and 100 nL reservoir solution (45% v/v 2-methyl-2,4-pentanediol (MPD), 0.2 M calcium chloride, 0.1 M Bis-Tris at pH 6.5). Crystals were cryoprotected with a multicomponent cryoprotectant (12,5% v/v diethylene glycol, 12.5% v/v MPD, 37,5% v/v 1,2-propanediol and 12,5% v/v dimethyl sulfoxide), harvested with 0.1-0.2 LithoLoops, and flash-vitrified in liquid nitrogen. X-ray diffraction datasets were collected at the XALOC (*28*) beamline of the ALBA synchrotron (Cerdanyola, Catalonia, Spain).

A diffraction dataset was collected at 2.8 A resolution and after data processing with Aimless (*29*) the structure was solved by molecular replacement with Phaser (*30*) using the original design model. The structures were refined using Refmac5 (*31*), Buster/TNT (*32*) and manual building with Coot (*33*).

### Tb3+ binding luminescence measurements

Time-resolved luminescence emission spectra and intensities were measured on a Synergy H1 hybrid multi-mode reader (BioTek) in flat bottom, black polystyrene, 96-well half-area microplates (Corning 3694). For Tb^3+^ luminescence experiments, protein samples were prepared in 20 mM Tris, 50 mM NaCl, pH 7.4. A TbCl_3_ (Sigma-Aldrich, 451304-1G) stock solution was prepared in the same protein buffer. Time-resolved luminescence intensities were measured using excitation wavelength λ_ex_ = 280 nm and emission wavelength λ_em_ = 544 nm with a delay of 300 ms, 1 ms collection time and 100 readings per data point. Time-resolved luminescence emission spectra between 500 nm and 600 nm were collected in 5 nm increments. For Tb^3+^ titrations, the collected time-resolved luminescence emission intensities at λ_em_ = 544 nm were normalized to obtain protein bound fractions, and the normalized data was fit to a 1:1 binding model using non-linear least squares regression. Ca^2+^ binding was probed by measuring the decrease of time-resolved terbium luminescence emission intensity at λ_em_ = 544 nm. CaCl_2_ was titrated in the same protein sample buffer into 10 μM protein and 10 μM Tb^3+^. The Ca^2+^ binding affinity constant (Kd(Ca^2+^)) was calculated from the apparent Kd (Kapp(Ca^2+^)) using the following equation: Kd(Ca^2+^) = Kapp(Ca^2+^) / (1+[Tb^3+^]/Kd(Tb^3+^)); where Kd(Tb^3+^) is the terbium-binding affinity constant calculated from Tb^3+^ titrations.

## ACKNOWLEDGMENTS

We thank F.X. Gomis-Rüth for help with crystallographic refinement. We also thank Joan Pous and Monica Buxaderas from the joint IBMB/IRB Automated Crystallography Platform and the Protein Purification Service for assistance during SEC-MALS, purification procedures, and crystallization experiments. E.M. acknowledges a Ramon y Cajal grant (RYC2018-025295-I) and two other grants from the Spanish Ministry of Science and Innovation (EUR2020-112164 and PID2020-120098GA-I00). J.R.T. was supported by an EMBO postdoctoral fellowship (under grant agreement ALTF 145-2021). X-ray diffraction data collection experiments were performed at the XALOC beamline at ALBA Synchrotron with the collaboration of ALBA staff. We acknowledge computing resources provided by the Galicia Supercomputing Center (CESGA).

## REFERENCES

1. J. Yélamos, “Current innovative engineered antibodies” in International Review of Cell and Molecular Biology (Elsevier, 2022; https://linkinghub.elsevier.com/retrieve/pii/S1937644822000284), vol. 369, pp. 1–43.

2. H. Kaplon, A. Chenoweth, S. Crescioli, J. M. Reichert, Antibodies to watch in 2022. mAbs. 14, 2014296 (2022).

3. C. Jost, A. Plückthun, Engineered proteins with desired specificity: DARPins, other alternative scaffolds and bispecific IgGs. Curr. Opin. Struct. Biol. 27, 102–112 (2014).

4. J. R. Kintzing, M. V. Filsinger Interrante, J. R. Cochran, Emerging Strategies for Developing Next-Generation Protein Therapeutics for Cancer Treatment. Trends Pharmacol. Sci. 37, 993–1008 (2016).

5. F. Sha, G. Salzman, A. Gupta, S. Koide, Monobodies and other synthetic binding proteins for expanding protein science. Protein Sci. 26, 910–924 (2017).

6. R. E. Bird, K. D. Hardman, J. W. Jacobson, S. Johnson, B. M. Kaufman, S.-M. Lee, T. Lee, S. H. Pope, G. S. Riordan, M. Whitlow, Single-Chain Antigen-Binding Proteins. Science. 242, 423–426 (1988).

7. X. Chen, J. L. Zaro, W.-C. Shen, Fusion protein linkers: Property, design and functionality. Adv. Drug Deliv. Rev. 65, 1357–1369 (2013).

8. J. S. Richardson, D. C. Richardson, Natural β-sheet proteins use negative design to avoid edge-to-edge aggregation. Proc. Natl. Acad. Sci. 99, 2754–2759 (2002).

9. T. M. Chidyausiku, S. R. Mendes, J. C. Klima, U. Eckhard, S. Houliston, M. Nadal, J. Roel-Touris, T. Guevara, H. K. Haddox, A. Moyer, C. H. Arrowsmith, F. X. Gomis-Rüth, D. Baker, E. Marcos, “De Novo Design of Immunoglobulin-like Domains” (preprint, Biochemistry, 2021), doi:10.1101/2021.12.20.472081.

10. J. K. Leman, B. D. Weitzner, S. M. Lewis, J. Adolf-Bryfogle, N. Alam, R. F. Alford, M. Aprahamian, D. Baker, K. A. Barlow, P. Barth, B. Basanta, B. J. Bender, K. Blacklock, J. Bonet, S. E. Boyken, P. Bradley, C. Bystroff, P. Conway, S. Cooper, B. E. Correia, B. Coventry, R. Das, R. M. De Jong, F. DiMaio, L. Dsilva, R. Dunbrack, A. S. Ford, B. Frenz, D. Y. Fu, C. Geniesse, L. Goldschmidt, R. Gowthaman, J. J. Gray, D. Gront, S. Guffy, S. Horowitz, P.-S. Huang, T. Huber, T. M. Jacobs, J. R. Jeliazkov, D. K. Johnson, K. Kappel, J. Karanicolas, H. Khakzad, K. R. Khar, S. D. Khare, F. Khatib, A. Khramushin, I. C. King, R. Kleffner, B. Koepnick, T. Kortemme, G. Kuenze, B. Kuhlman, D. Kuroda, J. W. Labonte, J. K. Lai, G. Lapidoth, A. Leaver-Fay, S. Lindert, T. Linsky, N. London, J. H. Lubin, S. Lyskov, J. Maguire, L. Malmström, E. Marcos, O. Marcu, N. A. Marze, J. Meiler, R. Moretti, V. K. Mulligan, S. Nerli, C. Norn, S. Ó’Conchúir, N. Ollikainen, S. Ovchinnikov, M. S. Pacella, X. Pan, H. Park, R. E. Pavlovicz, M. Pethe, B. G. Pierce, K. B. Pilla, B. Raveh, P. D. Renfrew, S. S. R. Burman, A. Rubenstein, M. F. Sauer, A. Scheck, W. Schief, O. Schueler-Furman, Y. Sedan, A. M. Sevy, N. G. Sgourakis, L. Shi, J. B. Siegel, D.-A. Silva, S. Smith, Y. Song, A. Stein, M. Szegedy, F. D. Teets, S. B. Thyme, R. Y.-R. Wang, A. Watkins, L. Zimmerman, R. Bonneau, Macromolecular modeling and design in Rosetta: recent methods and frameworks. Nat. Methods. 17, 665–680 (2020).

11. J. Jumper, R. Evans, A. Pritzel, T. Green, M. Figurnov, O. Ronneberger, K. Tunyasuvunakool, R. Bates, A. Žídek, A. Potapenko, A. Bridgland, C. Meyer, S. A. A. Kohl, A. J. Ballard, A. Cowie, B. Romera-Paredes, S. Nikolov, R. Jain, J. Adler, T. Back, S. Petersen, D. Reiman, E. Clancy, M. Zielinski, M. Steinegger, M. Pacholska, T. Berghammer, S. Bodenstein, D. Silver, O. Vinyals, A. W. Senior, K. Kavukcuoglu, P. Kohli, D. Hassabis, Highly accurate protein structure prediction with AlphaFold. Nature. 596, 583–589 (2021).

12. K. Tunyasuvunakool, J. Adler, Z. Wu, T. Green, M. Zielinski, A. Žídek, A. Bridgland, A. Cowie, C. Meyer, A. Laydon, S. Velankar, G. J. Kleywegt, A. Bateman, R. Evans, A. Pritzel, M. Figurnov, O. Ronneberger, R. Bates, S. A. A. Kohl, A. Potapenko, A. J. Ballard, B. Romera-Paredes, S. Nikolov, R. Jain, E. Clancy, D. Reiman, S. Petersen, A. W. Senior, K. Kavukcuoglu, E. Birney, P. Kohli, J. Jumper, D. Hassabis, Highly accurate protein structure prediction for the human proteome. Nature. 596, 590–596 (2021).

13. Y. Zhang, Skolnick, J., TM-align: a protein structure alignment algorithm based on the TM-score. Nucleic Acids Res. 33, 2302–2309 (2005).

14. E. Marcos, T. M. Chidyausiku, A. C. McShan, T. Evangelidis, S. Nerli, L. Carter, L. G. Nivón, A. Davis, G. Oberdorfer, K. Tripsianes, N. G. Sgourakis, D. Baker, De novo design of a non-local β-sheet protein with high stability and accuracy. Nat. Struct. Mol. Biol. 25, 1028–1034 (2018).

15. M. Siedlecka, G. Goch, A. Ejchart, H. Sticht, A. Bierzynski, Alpha-helix nucleation by a calcium-binding peptide loop. Proc. Natl. Acad. Sci. 96, 903–908 (1999).

16. P.-S. Huang, Y.-E. A. Ban, F. Richter, I. Andre, R. Vernon, W. R. Schief, D. Baker, RosettaRemodel: A Generalized Framework for Flexible Backbone Protein Design. PLoS ONE. 6, e24109 (2011).

17. J. C. Klima, L. A. Doyle, J. D. Lee, M. Rappleye, L. A. Gagnon, M. Y. Lee, E. P. Barros, A. A. Vorobieva, J. Dou, S. Bremner, J. S. Quon, C. M. Chow, L. Carter, D. L. Mack, R. E. Amaro, J. C. Vaughan, A. Berndt, B. L. Stoddard, D. Baker, Incorporation of sensing modalities into de novo designed fluorescence-activating proteins. Nat. Commun. 12, 856 (2021).

18. P. Bradley, K. M. S. Misura, D. Baker, Toward High-Resolution de Novo Structure Prediction for Small Proteins. Science. 309, 1868–1871 (2005).

19. E. Marcos, D. Silva, Essentials of de novo protein design: Methods and applications. WIREs Comput. Mol. Sci. 8 (2018), doi:10.1002/wcms.1374.

20. N. Koga, R. Tatsumi-Koga, G. Liu, R. Xiao, T. B. Acton, G. T. Montelione, D. Baker, Principles for designing ideal protein structures. Nature. 491, 222–227 (2012).

21. S. J. Fleishman, A. Leaver-Fay, J. E. Corn, E.-M. Strauch, S. D. Khare, N. Koga, J. Ashworth, P. Murphy, F. Richter, G. Lemmon, J. Meiler, D. Baker, RosettaScripts: A Scripting Language Interface to the Rosetta Macromolecular Modeling Suite. PLoS ONE. 6, e20161 (2011).

22. M. Mirdita, K. Schütze, Y. Moriwaki, L. Heo, S. Ovchinnikov, M. Steinegger, ColabFold: making protein folding accessible to all. Nat. Methods. 19, 679–682 (2022).

23. M. Steinegger, J. Söding, MMseqs2 enables sensitive protein sequence searching for the analysis of massive data sets. Nat. Biotechnol. 35, 1026–1028 (2017).

24. D. Case et al., AMBER 14 (University of California, San Francisco, 2014).

25. J. A. Maier, C. Martinez, K. Kasavajhala, L. Wickstrom, K. E. Hauser, C. Simmerling, ff14SB: Improving the Accuracy of Protein Side Chain and Backbone Parameters from ff99SB. J. Chem. Theory Comput. 11, 3696–3713 (2015).

26. G. M. Bradbrook, T. Gleichmann, S. J. Harrop, J. Habash, J. Raftery, J. Kalb (Gilboa), J. Yariv, I. H. Hillier, J. R. Helliwell, X-Ray and molecular dynamics studies of concanavalin-A glucoside and mannoside complexes Relating structure to thermodynamics of binding. J. Chem. Soc. Faraday Trans. 94, 1603–1611 (1998).

27. M. J. Abraham, T. Murtola, R. Schulz, S. Páll, J. C. Smith, B. Hess, E. Lindahl, GROMACS: High performance molecular simulations through multi-level parallelism from laptops to supercomputers. SoftwareX. 1-2, 19–25 (2015).

28. J. Juanhuix, F. Gil-Ortiz, G. Cuní, C. Colldelram, J. Nicolás, J. Lidón, E. Boter, C. Ruget, S. Ferrer, J. Benach, Developments in optics and performance at BL13-XALOC, the macromolecular crystallography beamline at the Alba Synchrotron. J. Synchrotron Radiat. 21, 679–689 (2014).

29. P. R. Evans, G. N. Murshudov, How good are my data and what is the resolution? Acta Crystallogr. D Biol. Crystallogr. 69, 1204–1214 (2013).

30. A. J. McCoy, R. W. Grosse-Kunstleve, P. D. Adams, M. D. Winn, L. C. Storoni, R. J. Read, Phaser crystallographic software. J. Appl. Crystallogr. 40, 658–674 (2007).

31. G. N. Murshudov, P. Skubák, A. A. Lebedev, N. S. Pannu, R. A. Steiner, R. A. Nicholls, M. D. Winn, F. Long, A. A. Vagin, REFMAC5 for the refinement of macromolecular crystal structures. Acta Crystallogr. D Biol. Crystallogr. 67, 355–367 (2011).

32. BUSTER version 2.10 (Global Phasing Ltd., Cambridge (UK) (2017).

33. A. Casañal, B. Lohkamp, P. Emsley, Current developments in Coot for macromolecular model building of Electron Cryo-microscopy and Crystallographic Data. Protein Sci. 29, 1055–1064 (2020).

