## Supplementary Information for "Design of extended metal-binding β-sandwiches from *de novo* immunoglobulin domains"

*for*

**Table S1. Crystallographic data.**

| *Dataset* | **dIG14-scdim** |
|---|---|
| Beam line (synchrotron) | XALOC (ALBA) |
| Space group / complexes per a.u. [a] | $P4_32_12$ / 2 |
| Cell constants (a,b and c in Å) | 72.01, 72.01, 97.07 |
| Wavelength (Å) | 0.97926 |
| Measurements / unique reflections | 76,734 / 6749 |
| Resolution range (Å) (outermost shell) [c] | 48.53-2.80 / (2.95-2.8) |
| Completeness (%) / $R_{merge}$ | 99.9 (99.9) / 0.094 (3.698) |
| $R_{pim}$ / CC($^1/_2$) | 0.029 (1.104) / 1 (0.585) |
| Average intensity [d] | 16.2 (0.7) |
| B-Factor (Wilson) (Å$^2$) / Aver. multiplicity | 98.2 / 11.4 (12.1) |
| Resolution range used for refinement (Å) | 45.09 – 2.80 |
| Reflections used (test set) | 6712 (678) |
| Crystallographic $R_{factor}$ (free $R_{factor}$) | 0.225 (0.284) |
| Non-H protein atoms/waters/ligands per a.u. | 1137 / 17 / 6 |
| *Rmsd* from target values | |
| bonds (Å) / angles (°) | 0.013 / 1.25 |
| Average B-factor (Å$^2$) | 104.0 |
| Protein contacts and geometry analysis [b] | |
| Ramachandran favoured / outliers / all analysed | 130 (94%) / 2 / 139 |
| Bond-length / bond-angle / chirality / planarity outliers | 0 / 0 / 0 / 0 |
| Side-chain outliers | 12 (9%) |
| All-atom clashes / clashscore [b] | 20 / 9 |
| RSRZ[a] outliers / $F_o$:$F_c$ correlation | 1 (0.709 %) [b] / 0.94 |
| PDB access code | XXXX |

[a] Abbreviations: a.u., asymmetric unit; RSRZ, real-space R-value Z-score.

[b] According to the wwPDB Validation Service (https://wwpdb-validation.wwpdb.org/validservice). [c] Values in parenthesis refer to the outermost resolution shell if not otherwise indicated.

[d] Average intensity is <I/σ(I)> of unique reflections after merging.

**Table S2. Structural similarity metrics and confidence values of AlphaFold2 predictions for each of our selected mono-functionalized designs.** RMSDs are calculated on Cα atoms after superimposition.

*model062_2_EF_1*

| Prediction | RMSD (Å) | | | pLDDT | | |
|---|---|---|---|---|---|---|
| | Total | Scaffold | Motif | Total | Scaffold | Motif |
| *Rank 1* | 0.97 | 0.18 | 1.17 | 96.3 | 96.5 | 93.7 |
| *Rank 2* | 0.97 | 0.21 | 1.16 | 95.9 | 96.1 | 93.5 |
| *Rank 3* | 0.91 | 0.16 | 1.14 | 95.8 | 96.1 | 92.5 |
| *Rank 4* | 0.99 | 0.17 | 1.14 | 95.1 | 95.5 | 91.4 |
| *Rank 5* | 0.81 | 0.18 | 1.15 | 95.0 | 95.5 | 90.2 |

*model004_2_EF_6*

| Prediction | RMSD (Å) | | | pLDDT | | |
|---|---|---|---|---|---|---|
| | Total | Scaffold | Motif | Total | Scaffold | Motif |
| *Rank 1* | 0.87 | 0.25 | 0.9 | 94.0 | 94.9 | 86.8 |
| *Rank 2* | 0.78 | 0.24 | 0.89 | 92.1 | 93.2 | 83.1 |
| *Rank 3* | 0.79 | 0.21 | 0.88 | 92.0 | 93.4 | 80.1 |
| *Rank 4* | 1.20 | 0.23 | 0.88 | 90.4 | 92.8 | 71.2 |
| *Rank 5* | 2.68 | 1.88 | 1.03 | 89.9 | 91.4 | 77.8 |

*model015_2_EF_2*

| Prediction | RMSD (Å) | | | pLDDT | | |
|---|---|---|---|---|---|---|
| | Total | Scaffold | Motif | Total | Scaffold | Motif |
| *Rank 1* | 0.82 | 0.19 | 1.26 | 92.1 | 94.3 | 77.3 |
| *Rank 2* | 0.85 | 0.26 | 1.40 | 91.4 | 92.4 | 84.6 |
| *Rank 3* | 0.78 | 0.26 | 1.22 | 91.2 | 93.2 | 76.9 |
| *Rank 4* | 1.32 | 0.21 | 1.40 | 90.7 | 93.0 | 74.5 |
| *Rank 5* | 2.42 | 1.76 | 1.71 | 87.1 | 89.1 | 73.3 |

*model018_2_EF_7*

| Prediction | RMSD (Å) | | | pLDDT | | |
|---|---|---|---|---|---|---|
| | Total | Scaffold | Motif | Total | Scaffold | Motif |
| *Rank 1* | 0.75 | 0.23 | 1.16 | 93.4 | 94.2 | 88.2 |
| *Rank 2* | 0.66 | 0.20 | 1.06 | 92.6 | 93.7 | 85.6 |
| *Rank 3* | 0.82 | 0.34 | 1.18 | 92.0 | 93.1 | 85.0 |
| *Rank 4* | 0.89 | 0.39 | 1.12 | 91.5 | 92.6 | 84.9 |
| *Rank 5* | 2.03 | 1.41 | 1.22 | 77.9 | 78.7 | 72.5 |

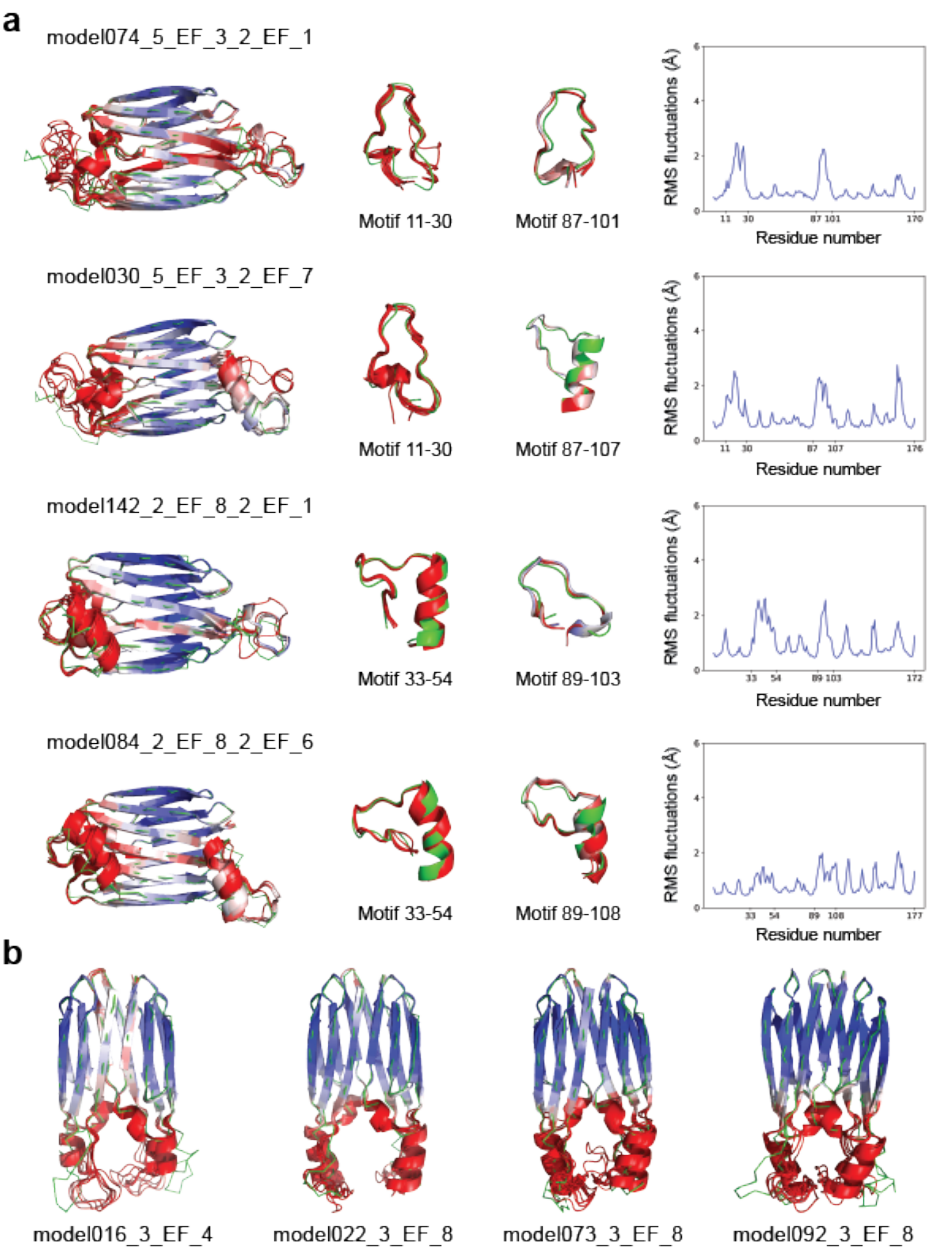

**Fig. S1. Computational design and characterization of dIG14-scdim with two EF-hand motifs. a,** From left to right: Selected designs with functional metal-binding loops at either side of the scaffold (green) with their corresponding AF2 predictions colored according to pLDDT (scale 70 (red) to 100 (blue)), closer look at the inserted motifs and Root Mean Square fluctuations obtained from molecular dynamics simulations. X-axis shows the grafted motifs and the last residue of the scaffold. Y-axis indicates fluctuations in Angstroms for each residue. **b,** Selected designs with functional metal-binding loops at the same side of the scaffold (green) with their corresponding AF2 predictions colored by increasing in pLDDT (scale 70 to 100).

**Table S3. Structural similarity metrics and confidence values of AlphaFold2 predictions for each of our selected bi-functionalized designs.** RMSDs are calculated on Cα atoms after superimposition.

*model074_5_EF_3_2_EF_1*

| Prediction | RMSD (Å) | | | pLDDT |
|---|---|---|---|---|
| | Scaffold | Motif 11-30 | Motif 87-101 | |
| *Rank 1* | 0.54 | 5.42 | 3.49 | 81.1 |
| *Rank 2* | 0.58 | 7.53 | 2.12 | 80.3 |
| *Rank 3* | 1.17 | 5.89 | 3.14 | 78.6 |
| *Rank 4* | 1.30 | 8.15 | 1.54 | 76.3 |
| *Rank 5* | 0.85 | 8.73 | 4.98 | 69.1 |

*model030_5_EF_3_2_EF_7*

| Prediction | RMSD (Å) | | | pLDDT |
|---|---|---|---|---|
| | Scaffold | Motif 11-30 | Motif 87-107 | |
| *Rank 1* | 0.47 | 5.63 | 2.51 | 88.6 |
| *Rank 2* | 0.40 | 4.79 | 1.86 | 83.6 |
| *Rank 3* | 0.54 | 8.28 | 2.49 | 80.2 |
| *Rank 4* | 0.38 | 4.83 | 12.33 | 79.0 |
| *Rank 5* | 0.51 | 5.04 | 1.97 | 76.0 |

*model142_2_EF_8_2_EF_1*

| Prediction | RMSD (Å) | | | pLDDT |
|---|---|---|---|---|
| | Scaffold | Motif 33-54 | Motif 89-103 | |
| *Rank 1* | 0.21 | 2.91 | 3.51 | 88.5 |
| *Rank 2* | 0.32 | 3.87 | 3.24 | 87.9 |
| *Rank 3* | 0.27 | 3.52 | 3.35 | 87.0 |
| *Rank 4* | 0.42 | 3.96 | 6.42 | 82.02 |
| *Rank 5* | 0.42 | 3.75 | 2.68 | 79.7 |

*model084_2_EF_8_2_EF_6*

| Prediction | RMSD (Å) | | | pLDDT |
|---|---|---|---|---|
| | Scaffold | Motif 33-54 | Motif 89-108 | |
| *Rank 1* | 0.28 | 3.59 | 2.62 | 83.8 |
| *Rank 2* | 0.38 | 3.64 | 2.18 | 83.0 |
| *Rank 3* | 0.26 | 3.57 | 2.42 | 82.4 |
| *Rank 4* | 0.49 | 3.56 | 2.42 | 79.2 |
| *Rank 5* | 0.40 | 3.47 | 3.04 | 78.7 |

*model016_3_EF_4*

| Prediction | RMSD (Å) | | | pLDDT |
|---|---|---|---|---|
| | Scaffold | Motif 11-29 | Motif 99-117 | |
| *Rank 1* | 0.68 | 4.89 | 5.94 | 85.8 |
| *Rank 2* | 0.66 | 5.04 | 5.71 | 83.9 |
| *Rank 3* | 0.74 | 5.46 | 5.92 | 82.8 |
| *Rank 4* | 0.76 | 5.86 | 7.00 | 80.0 |
| *Rank 5* | 0.99 | 6.66 | 8.17 | 79.6 |

*model022_3_EF_8*

| Prediction | RMSD (Å) | | | pLDDT |
|---|---|---|---|---|
| | Scaffold | Motif 11-33 | Motif 103-121 | |
| *Rank 1* | 1.00 | 5.19 | 5.28 | 84.3 |
| *Rank 2* | 1.02 | 3.78 | 4.31 | 83.8 |
| *Rank 3* | 0.95 | 3.84 | 4.33 | 82.7 |
| *Rank 4* | 0.94 | 3.71 | 4.22 | 82.4 |
| *Rank 5* | 1.00 | 4.87 | 5.42 | 82.3 |

*model073_3_EF_8*

| Prediction | RMSD (Å) | | | pLDDT |
|---|---|---|---|---|
| | Scaffold | Motif 11-33 | Motif 103-121 | |
| *Rank 1* | 0.66 | 2.75 | 2.31 | 83.8 |
| *Rank 2* | 0.64 | 2.92 | 2.73 | 81.5 |
| *Rank 3* | 0.82 | 3.78 | 3.28 | 80.3 |
| *Rank 4* | 1.15 | 6.07 | 6.06 | 80.3 |
| *Rank 5* | 1.08 | 3.70 | 3.26 | 78.6 |

*model092_3_EF_8*

| Prediction | RMSD (Å) | | | pLDDT |
|---|---|---|---|---|
| | Scaffold | Motif 11-33 | Motif 103-121 | |
| *Rank 1* | 1.20 | 6.62 | 7.02 | 85.1 |
| *Rank 2* | 1.27 | 6.56 | 6.46 | 82.3 |
| *Rank 3* | 0.85 | 7.22 | 6.90 | 81.5 |
| *Rank 4* | 1.26 | 6.16 | 7.30 | 81.5 |
| *Rank 5* | 0.99 | 5.69 | 4.55 | 78.7 |
